# Reversible temporally-specific inhibition of muscle using a light-activated chloride channel

**DOI:** 10.1101/398156

**Authors:** Christopher J. Gorini, Shrivats M. Iyer, Saurabh Vyas, Charu Ramakrishnan, Karl Deisseroth, Scott L. Delp

**Author notes:** Corresponding Author: Scott L. Delp; Clark Center S321, 318 Campus Drive, Stanford, CA 94305.

## Abstract

Achieving reversible, temporally specific inhibition of motor neurons has the potential to revolutionize treatment of disorders marked by muscle hyperactivity. Current treatment strategies are inadequate – surgical interventions are irreversible and pharmaceutical interventions have off-target effects. Optogenetic strategies for inhibiting muscle activity have theoretical promise; however, trafficking and expression problems have prevented translatable optogenetic suppression of muscle activity. Here, we exploit recent innovations in opsin engineering to demonstrate virally mediated, temporally specific optogenetic inhibition of motor neurons and muscle activity *in vivo*. We show that intra-muscular injection of adeno-associated virus serotype 6 can drive expression of the inhibitory channelrhodopsin mutant iC++ in both immunodeficient and wild-type mice. Illumination of the sciatic nerve in wild-type mice resulted in 64.8% inhibition of evoked twitch force. Optical excitation during tetanic stimulation in wild-type mice resulted in 59.1% inhibition at 10 Hz and 55.4% at 25 Hz. The extent of optogenetic inhibition was titratable, and ranged from 0% to 78.4% as illumination intensity was changed from 1 mW to 20 mW/mm^2^. These results could have therapeutic applicability to disorders such as spasticity, hypertonia and urinary incontinence, and provide a new tool to neuroscientists and muscle physiologists wishing to reversibly inhibit motor neuron activity *in vivo*.

## Introduction

Disorders of the neuromuscular system can compromise activities of daily living. While varying widely in their causes, many of these disorders have excessive muscle activity as a common symptom. This excessive activity adversely affects quality of life in different ways. Spasticity, for example, results in involuntary muscle contractions that interfere with precise movement and locomotion[1, 2]. In overactive bladder syndrome, excessive activation of the detrusor muscles of the bladder results in unwanted micturition[3, 4].

Surgical and pharmacological interventions to reduce muscle hyperactivity have important limitations. Surgical approaches include division of nerves through rhizotomy or peripheral neurectomy, which is irreversible and may cause unwanted results[5, 6]. Intrathecally administered baclofen is commonly used to diminish spasticity but does not inhibit specific muscles, and has implant associated morbidity[7]. Intramuscular injection of botulinum toxin has only transient effects, and is complicated by redosing induced tolerance[8]. Off-label use of medications such as diazepam and riluzole may offer some relief; however, they have unreliable efficacy and side effects such as fatigue, dizziness, depression and cognitive dullness[9, 10].

Optogenetic techniques enable temporally precise, reversible control of neuronal activity[11]. However, optogenetic strategies to excite neural activity have been more tractable than optogenetic silencing[12-16]. In our experience, this has proven particularly true in control of neuromuscular circuits[17]. Our laboratory, and others, have previously reported optogenetic *excitation* of muscle activity *in vivo*, using a transgenic approach and using viral or cell-transfer models that have some translational resonance[18, 19]. While we have been able to achieve optogenetic inhibition of muscle activity using a transgenic mouse line[18], the critical step of optogenetically inhibiting muscle through a translationally relevant approach has, until now, remained intractable. Here, we exploit recent innovations in opsin engineering[20] to show that the improved inhibitory opsin mutant, iC++, when virally delivered to peripheral motor neurons, enables inhibition of muscle activity *in vivo*.

## Materials and Methods

### Animal test subjects

All animal procedures were approved by the Administrative Panel on Laboratory Animal Care at Stanford University, in accordance with the “Guide for the Care and Use of Laboratory Animals” from the National Institutes of Health.

### Viral vectors

AAV6 expressing iC++ (AAV6-iC++) was produced at the University of North Carolina Vector Core Facility. The expression cassette comprised iC++ fused to yellow fluorescent protein (eYFP) under control of the human synapsin promoter. Genomic DNA was packaged into the AAV6 capsid using helper plasmids. A single viral batch was used with a dot-blot hybridization titer of 1.4×10^13^ v/ml.

### Intra-muscular viral injections

Healthy juvenile female C57BL/6 and *Foxn1*^*NU*^ (Nu/J) mice, aged 4 weeks (average weight 23.0 g), were obtained from Jackson Laboratories (Bar Harbor, Maine) and kept on a 12:12 light:dark cycle. Mice were anesthetized using isoflurane (1-3%) and the gastrocnemius was exposed. Using a syringe pump (World Precision Instruments, Sarasota, FL) fitted with a Hamilton syringe (Hamilton, Reno, NV), 12 µl of AAV6-hSyn-iC++-eYFP was injected into the medial and lateral left gastrocnemius across 4 sites, using a 35G needle, at a rate of 1.5 µl/min. The skin was then sutured closed and mice were allowed to recover. One week prior to euthanasia, using cervical dislocation under isoflurane, mice were injected with 12 µl of 4% Fluorogold (Santa Cruz Biotechnology, Dallas, TX) in the virus-injected muscle, using the procedures described previously[17, 18].

### Histology

Spinal cord and sciatic nerve, were dissected, post-fixed (4°C overnight), and transferred to 30% sucrose in PBS (4°C until sectioning). Tissues were embedded in Tissue-Tek OCT compound (Sakura, Holland) and sectioned on a Leica CM3050 cryostat (Leica, Buffalo Grove, IL). Spinal cords were cut in 40 µm longitudinal sections and 20 µm cross sections and mounted onto glass slides. Slides were stained and mounted in PVA-DABCO (Sigma-Aldrich, St. Louis, MO). Sciatic nerves were processed in the same manner with 20 µm section thickness. The percentage of Fluorogold positive motor neurons expressing iC++ was quantified in 13 out of 40 sections.

### *In vivo* measurements

Nu/J and wild-type mice were anesthetized with isoflurane (1–3%) and kept warm on an electrical heating pad (37°C). The hindlimb was shaved, and the sciatic nerve exposed, and the Achilles tendon isolated. The calcaneus was cut, and a small bone piece at the tendon was attached through a light-weight, rigid hook to a force transducer (0.3 mN resolution; 300C-LR; Aurora Scientific, Aurora, Ontario). Electromyography (EMG) electrodes from stainless steel, tetrafluoroethylene-coated wires (790700; A-M Systems, Carlsborg, Washington) were inserted into the medial gastrocnemius muscle belly and muscle–tendon junction with a ground electrode at the forelimb wrist.

A custom-built electrical cuff (Microprobes, Gaithersburg, MD) was placed around the sciatic nerve and motor nerves were electrically stimulated (Powerlab 16/35, ADinstruments, Colorado Springs, CO). Supramaximal stimulation was determined for each mouse by incrementally increasing voltage until twitch force amplitude plateaued. Twitch stimulation was 2 ms pulses at 1 Hz for 60-90 s at 50% supramaximal twitch force. Tetanic stimulation was 2 ms pulse width at 10 and 25 Hz for 60-90 s at approximately 50% of supramaximal twitch force. Rest periods were 120 s between trials.

### Illumination

Continuous illumination at the nerve was from a blue laser (473 nm, OEM Laser Systems, East Lansing MI) via a multimode optical fiber (365 µm diameter, 0.37 numerical aperture, Thorlabs, Newton NJ). Light power density was calculated as the power at the fiber tip measured with a power meter (PM100, Thorlabs) divided by the light spot area of 1 mm^2^ illuminating the nerve (light spot diameter of approximately 1.1 mm). Experiments with Nu/J and wild-type mice involved 2 to 4 hours of repeated electrical and optical stimulation (up to 75 trials per mouse). We observed no localized nerve damage and no systematic non-optogenetic change in force or EMG during this time. Each trial included consecutively 10 s pre-light, 30 s with light, and 20 s post-light twitches or 10 s pre-light, 70 s with light, and 20 s post-light twitches. The start, end, frequency, and pulse duration of the light within each trial was controlled using Powerlab 16/35 (ADinstruments, Colorado Springs, CO). To achieve optical inhibition, we illuminated axons in the nerve at the electrical stimulation cuff.

### Data analysis

Force and EMG data were recorded at a sampling frequency of 1 kHz using Powerlab 16/35 and analyzed using both Labchart 7.0 (ADinstruments, Colorado Springs, CO) and custom scripts written in MATLAB (Mathworks, Natick, MA). Passive force amplitude (muscle force with no activation) was measured for each trial prior to electrical stimulation of the nerve. Analyses were done on the active (total - passive) force.

Electrically evoked forces were calculated at three time-points: ‘pre-light’, during ‘light’, and ‘recovery’. As electrically evoked forces were observed to decay in an approximately exponential manner, the extent of optogenetically evoked inhibition was calculated based on mean evoked forces during the final 20% of optical stimulation (at which point inhibition intensity had stabilized. Depending on the duration of illumination, the time-period included varied from 2–24 seconds. For consistency, ‘pre-light’ and ‘recovery’ forces were also calculated as the mean forces evoked during the final 20% of each time-period.

Time-constants were determined for inhibition ‘onset’ by fitting observed force peaks to an exponential function of the form 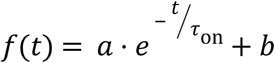.

### Statistics

Calculations were performed separately for each stimulation trial, and the inhibition parameters for all trials from a given mouse were then averaged together to determine the parameters for that mouse. Group data means and standard errors were then calculated as based on trial-averaged mouse results. P-values were calculated using the paired or unpaired Student’s *t*-test, as appropriate. Significance thresholds were set at P < 0.05 (*), P < 0.01 (**), and P < 0.001 (***). Effect sizes were calculated using the Hedges *g* metric, through the Measures of Effect Size Toolbox[21]. All grouped data are normalized and shown as mean ± s.e.m.

## Results

We began by conducting a broad-spectrum scan of adeno-associated viruses and inhibitory opsins to test whether different combinations of virus, opsin, dose and delivery route could produce sufficient opsin expression to enable silencing of muscle activity *in vivo*. Surprisingly, despite testing 23 such virus-opsin combinations *in vivo* (S1 Table), we were unable to achieve consistent, robust optogenetic inhibition of electrically evoked muscle twitches. Histological examination of the nerves and spinal cords of these mice indicated that the principal reason was a lack of sufficient expression and trafficking of the opsin transgene in primary motor neurons.

To overcome this, we adopted two new experimental approaches. First, because we had previously observed a strong immune response to peripheral adeno-associated virus administration[22], consistent with reports from clinical trials[23], we reasoned that performing our experiments in immunocompromised mice would be useful to validate our viral expression strategy. Second, because we had previously observed that channelrhodopsin-2 could be virally expressed with high efficiency in primary motor neurons[17], we reasoned that the recently developed inhibitory chloride-conducting channelrhodopsin mutant, iC++, may exhibit high transduction efficiency, and may therefore enable functional inhibition *in vivo*.

We assayed immunocompromised mice 5 weeks following injection of AAV6-hSyn-iC++-eYFP into the gastrocnemius muscle. In stark contrast to our previous inhibitory experiments, we observed that iC++ trafficked throughout the primary motor neuron with high efficacy (Fig 1a). We performed Fluorogold injections into the virally injected muscle to quantify the percentage of the motor pool transduced, and observed it to be ∼38% (Fig 1a, top, *n* = 2).

**Fig 1.**
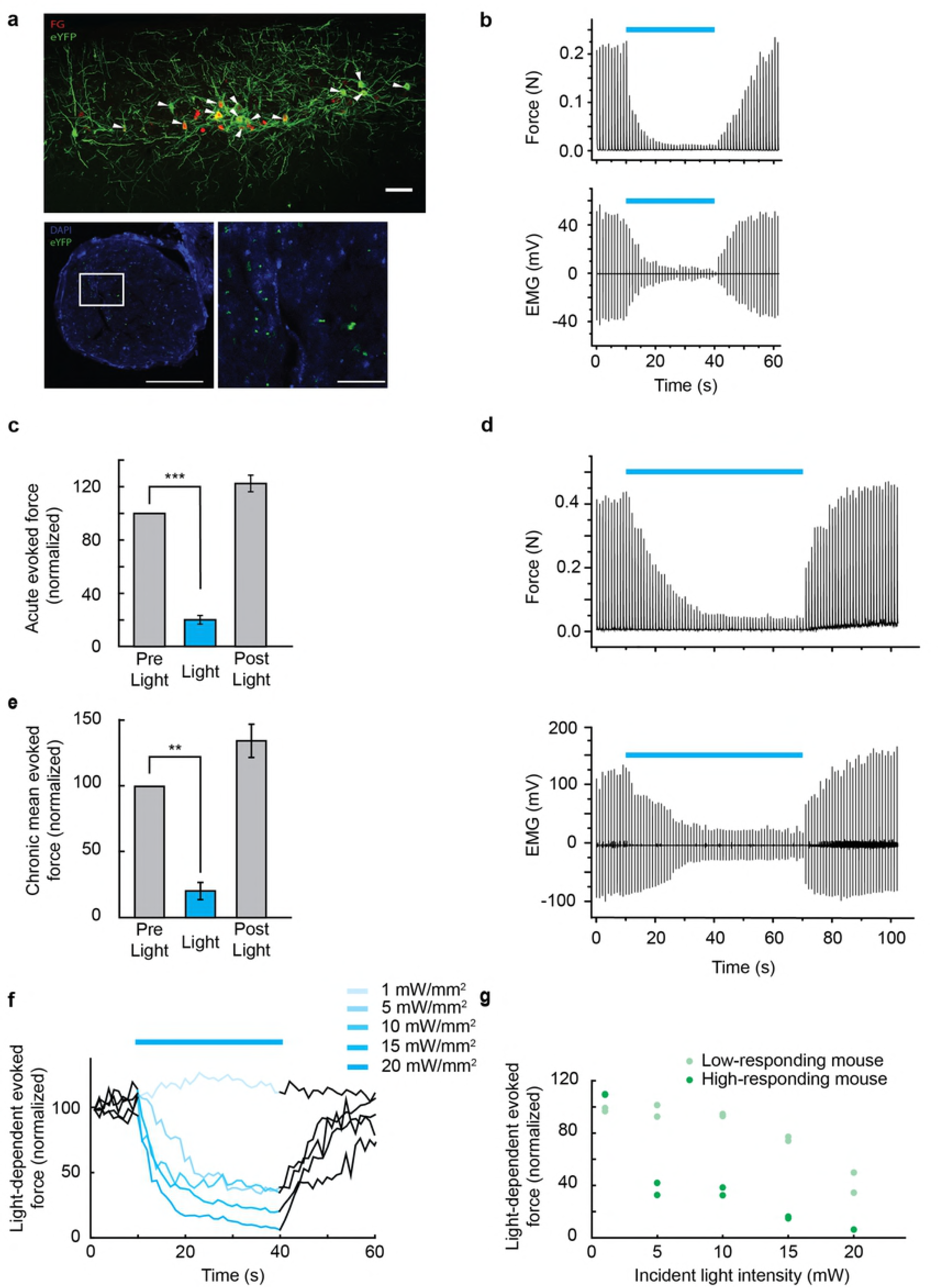
Reversible optogenetic inhibition of twitch force in muscle. a, top. Longitudinal section of lumbar spinal cord 5 weeks post intramuscular injection of iC++ (eYFP, green) and one week post fluorogold (FG, red) injection. Arrowheads denote co-localization of eYFP with fluorogold, representing an average of 38.1% of the total motor neuron population over 2 animals expressing iC++. Scale bar 100 µm. a, lower. Confocal image of a cross-section of the sciatic nerve showing motor nerve axons expressing eYFP from the fusion protein of iC++-eYFP (green) 5 weeks following intramuscular injection. Blue, DAPI, scale bar 100 µm. Right panel shows inset drawn in left panel. Scale bar 50 µm b. An example force trace is shown, top, with corresponding EMG, bottom. Electrical stimulation of the sciatic nerve of Nu/J mice (2ms pulse duration) was applied at 1Hz to produce a 50% maximal force for the duration of the 60s trial. Compared to pre-light force, we observed a 79.9 ± 3.2% average decrease in force occurring under constant blue light (473nm) illumination (20mW/mm^2^) for 30s, as indicated by blue bars (P = 8.1e^-09^, effect size = 4.56, n = 12). The mean time constant (**τ** onset) of the inhibition was calculated to be 4.08s ± 0.48 (r^2^ = 09.6 ± 0.006). c. Summary data from 12 animals representing the average inhibition at 1Hz electrically evoked force. d. Representative force trace of chronic inhibition (> 50s), top and corresponding EMG, bottom, with blue bars representing constant illumination of sciatic nerve. e. Summary bar graph of mean chronic force inhibition showing 75.4 ± 7.5% optical inhibition (P = 0.007, effect size = 2.50, n = 4) with a recovery of 133.2 ± 11.2% of pre-light force. All grouped data are normalized and shown as mean ± s.e.m. f. Twitch force inhibition was dependent on the density of illumination. Summary data of force traces at 50% maximum force (2 ms electrical pulse duration at 1 Hz) in response to varying light densities of blue light (20, 15, 10, 5, 1 mW/mm^2^, n=1), with maximum force inhibition occurring at 20mW/mm^2^. g. Example normalized plot of 4 mice (2 high-responding, dark green, and 2 low-responding, light green) representing the difference in light evoked inhibition over changes in light densities. Optically elicited force inhibition was shown to vary over the individual mice tested.

We then turned to functional assays of motor neuron inhibition. Following experimental methods described previously[18], we placed an electrical cuff around the sciatic nerve, and illuminated the region of the nerve that received electrical stimulation (constant light: 20 mW/mm^2^, 473 nm; electrical stimulation: 1 Hz, 2 ms), while simultaneously measuring force output and electromyographical activity (EMG) in the targeted muscle (S1 Fig). We observed that blue light (473 nm) illumination in iC++ injected mice was sufficient to block force generation in the targeted muscle, with concomitant reduction in the EMG waveform (Fig 1b). Importantly, optogenetically evoked inhibition was completely reversible, with force output returning to 122.6 ± 1.8 % (*n* = 12) of baseline pre-light force (Fig 1b,c). Across 12 mice, iC++ inhibition during electrically evoked twitch force was robust; at 1 Hz we observed an average inhibition of 79.9 ± 3.2% compared to pre-light baseline force (P = 8.10e^-09^, effect size = 4.6, Fig 1c). Interestingly, in contrast with previous reports of light-mediated inhibition in transgenic Thy1-NpHR2.0 mice[18], optogenetic inhibition and recovery with iC++ was not instantaneous, and followed a consistently exponential onset pattern (**τ**_on_ = 4.08s ± 0.49, *R*^*2*^ = 0.955, P = 0.006, Fig 1b). Inhibition could be sustained over a wide range of durations, lasting for as long as 70s, with no noticeable deleterious effects on post-light force production (Fig 1d,e). We did not observe any reduction in magnitude of inhibition over the course of the illumination period. The degree of optogenetic inhibition could be titrated over a range of 78.4% to 0% by changing illumination intensity from 20 mW/mm^2^ to 1 mW/mm^2^ (*n* = 1, Fig 1f), respectively. We observed that this intensity-dependence was a function of the animal’s light-sensitivity, and varied with the animals’ peak achievable inhibition. We categorized 4 animals, 2 high-responding (dark green) and 2 low-responding (light green) that best displayed this variation (*n* = 4, Fig 1g).

We next examined whether iC++ could suppress higher frequencies of electrically evoked muscle activity (10 Hz, 2 ms; 25 Hz, 2 ms). We observed significant optogenetic silencing of muscle force production during electrical stimulation at 10 Hz, with the force envelope decaying an average of 79.5 ± 9.2% (P= 0.009, effect size = 0.096, *n* = 4) of the immediate pre-light force levels (Fig 2a,b). At 25 Hz contraction, due to electrical stimulation-evoked fatigue, muscle force after illumination did not completely recover to peak pre-light levels; however, it did reach 66.9 ± 3.6% (*n* = 5) average recovery of initial pre-light force measurements (Fig 2c,d) following a 71.3 ± 11.4% (P = 0.001, effect size = 1.8) optical inhibition.

**Fig 2.**
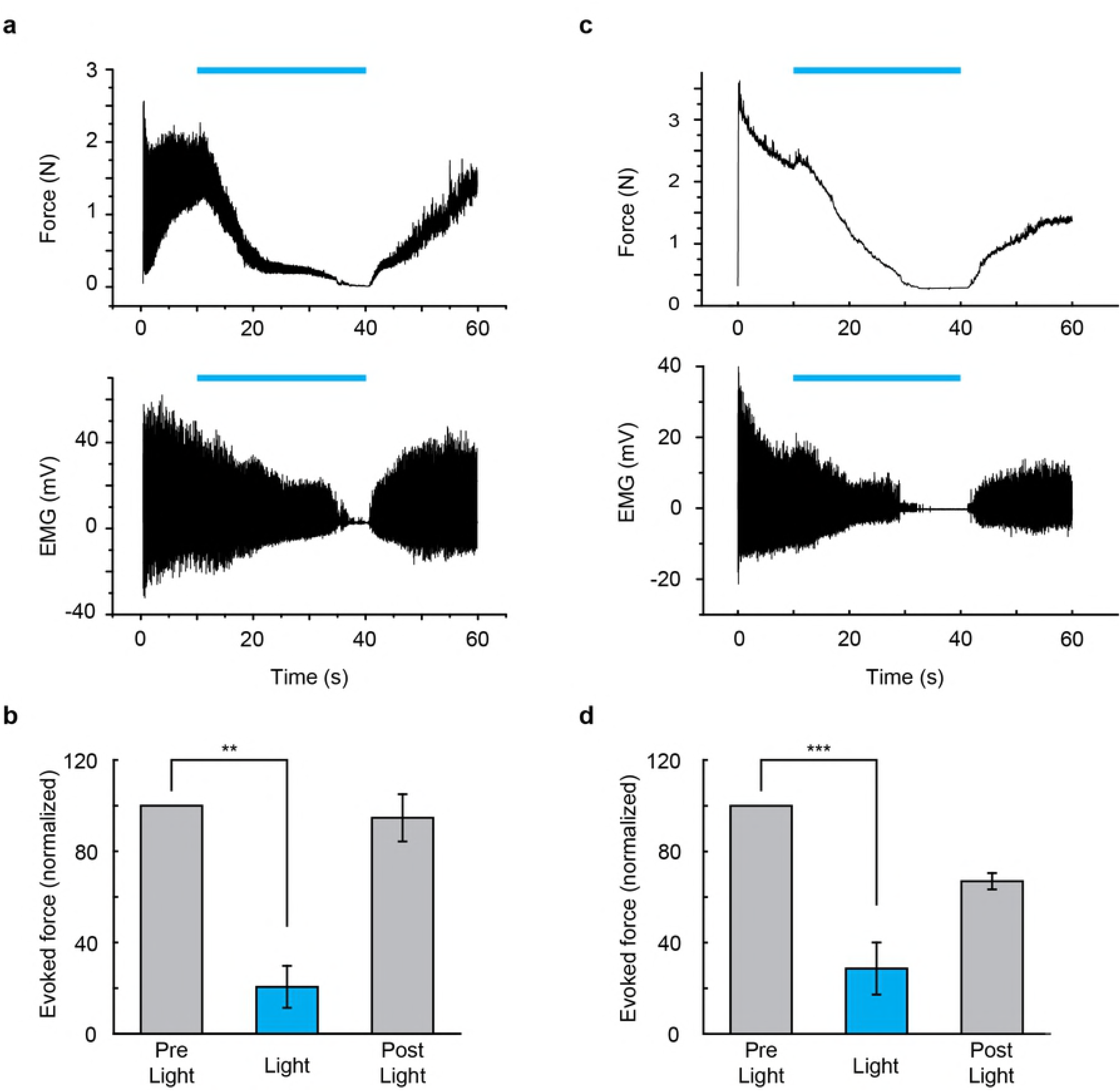
iC++ mediated tetanic force inhibition at 10 Hz and 25 Hz. a. Example force (top) and corresponding EMG (bottom) tetanic trace elicited by 60 s of 2 ms electrical stimulation at 10Hz with 30 s of iC++ mediated inhibition using 473 nm constant light (blue bars). b. 10 Hz summary data show an average optogenetic elicited inhibition of 79.5 ± 9.2% compared to pre-light levels with a 94.7 ± 10.4% recovery (P = 0.009, effect size = 0.957, n = 4). c. Representative trace of tetanic force (top) and EMG (bottom) generated by 2ms electrical stimulation at 25 Hz for 60 s. d. Summary data showing a mean inhibition of 71.3 ± 11.4% compared to pre-light levels (P = 0.001, effect size = 1.89, n=5). Average post-light force recovery was 66.9 ± 3.6% of baseline. All grouped data are normalized and shown as mean ± s.e.m.

Our initial experiments were performed in immunocompromised mice. In the clinical setting, gene therapy is often performed with simultaneous immunosuppression[23]; however, as this poses significant side-effects[24], we examined if we could achieve virally-mediated optogenetic inhibition in wild-type mice. We injected AAV6-hSyn-iC++-eYFP into the gastrocnemius muscle of 4 week old wild-type mice and, in contrast with our previous efforts with other opsin-virus cocktails, we observed robust expression in primary motor neurons in these mice 5 weeks post-injection (Fig 3a). Opsin expression patterns were broadly similar to those seen in immunocompromised mice.

**Fig 3.**
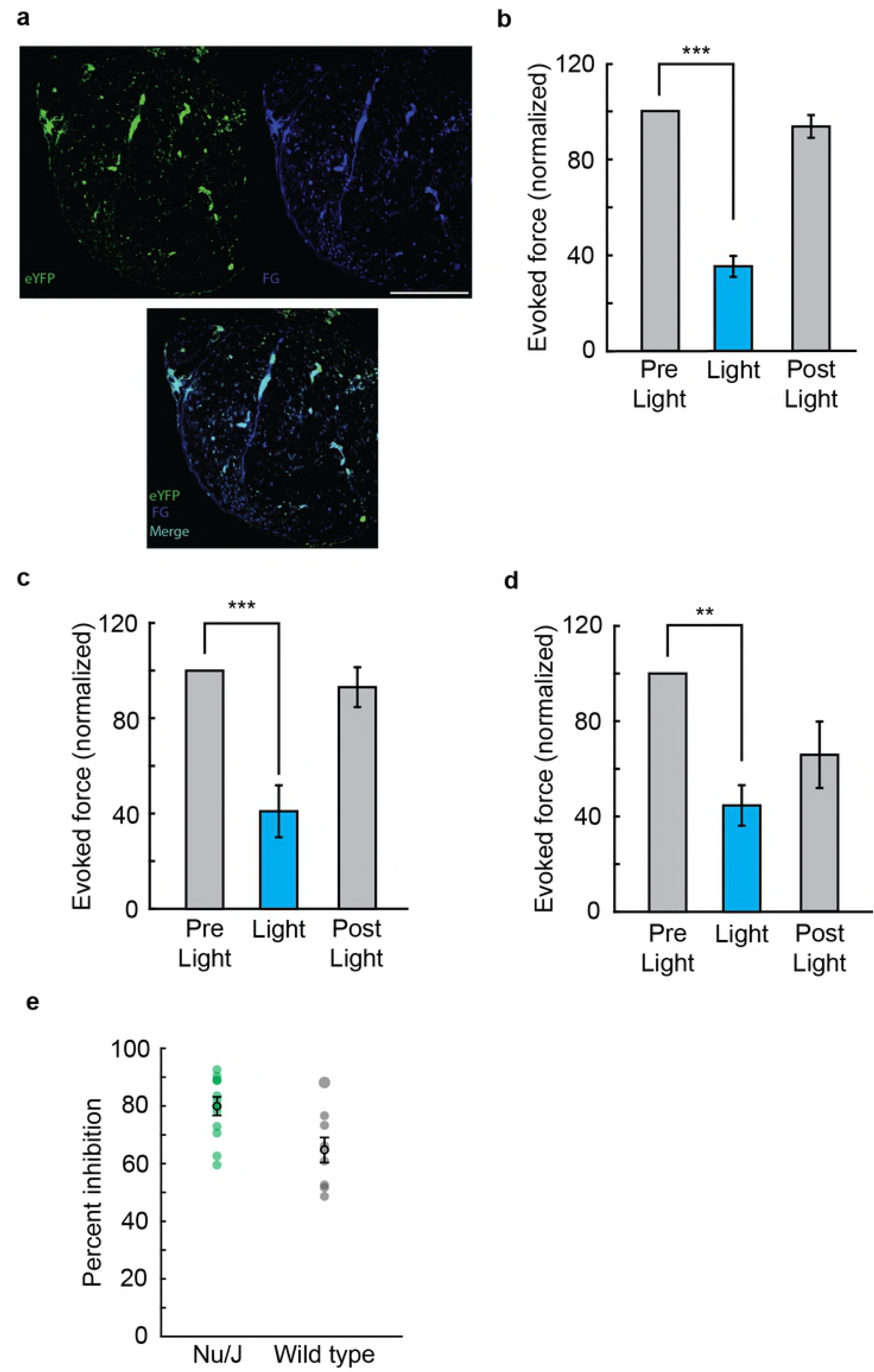
Optical inhibition of electrically evoked muscle force in wild type animals. a. Confocal image of sciatic nerve cross-section in wild type mouse expressing eYFP (top, left, green) and FG (top, right, blue) in motor axons 5 weeks post injection. FG was injected one-week prior to sacrifice. a, bottom. A merge of eYFP and FG expressing fiber. Scale bar 100 µm. b. Average iC++ mediated force inhibition over 9 wild type animals at 1 Hz was 64.8 ± 4.3% with a mean recovery of 93.5 ± 1.6% (P = 0.0004, effect size = 2.33, **τ** onset = 7.4s ± 0.8, r^2^ = 0.92 ±0.01). c,d. Wild type muscle inhibition from 10 and 25 Hz recordings, respectively, was similar to nude mice data. c. At 10 Hz evoked force, average inhibition observed was 59.1 ± 10.9% with a 93.0 ± 8.3% recovery of pre-light force (P = 0.005, effect size = 1.35, n= 6). d. The mean electrically evoked force at 25 Hz was optically suppressed by 55.4 ± 8.5% and exhibited a 65.7 ± 13.9% recovery (P = 0.012, effect size = 3.76, n= 4). e. Scatter plot representing the individual distribution of mean force inhibition across nude (green, average = 79.9 ± 3.2%, P = 8.1e^-09^, n=12) and wild type animals (gray, average = 64.8 ± 4.3%, P = 0.0004, n=9) at 1 Hz. All grouped data are shown as mean ± s.e.m.

We repeated all previously described *in vivo* experiments, achieving a substantially similar degree of inhibition (Fig 3b-d). We obtained significant optogenetic inhibition at 1 Hz twitch force with high efficacy (average suppression of 64.8 ± 4.3%, P = 0.0004, effect size = 2.33, *n* = 9, Fig 3b) and were able to inhibit tetanic force production in wild-type mice (Fig 3c,d). The extent of inhibition achievable in wild-type mice (gray) was comparable to that observed in immunocompromised mice (green, Fig 3e).

## Discussion

Here, we report light-mediated inhibition of peripheral motor neurons in nontransgenic mice. This proved exceptionally difficult to achieve (S1 Table). Clinical translation of optogenetic methods to inhibit muscle will require several additional innovations. Chief among these will be the development of improved viral vectors with greater transduction efficacy and reduced immunogenic profile. Additional improvements in inhibitory opsin photocurrents are also likely to be of use[25].

A benefit of the optogenetic approach described here is that it allows inhibition of motor neurons through illumination of the peripheral axon, without requiring illumination of the soma within the spinal cord. In eventual clinical translation, it is possible to imagine illuminating peripheral axons through the use of a wirelessly powered nerve cuff[26]. Key remaining hurdles to translation include establishing inhibitory control of motor neurons in species other than mouse and demonstrating optogenetic silencing of muscle during animal locomotion. Additional translational targets are likely to include the urinary system in disorders such as urinary incontinence and detrusor-sphincter dyssynergia. Clinical modulation here would involve injecting the relevant muscles, such as the external urethral sphincter, with retrogradely transducing AAVs, and then illuminating the transduced efferent nerves to induce micturition on command[27-29].

The use of a novel light-activated chloride mediated inhibitory channel may also be of interest in cardiac resynchronization therapy. To date, optogenetic inhibition of cardiomyoctes has been limited to cultured rat cardiomyocte cells transfected with the inhibitory proton pumps, Arch3[30] or Arch-T[31]. Although suppression of electrical activity was observed, no *in vivo* aspects of function were tested. The light-activated chloride channel, iC++, may prove useful in inhibition of cardiac muscle, *in vivo,* to achieve a tunable, temporally precise inhibition for cardiac abnormalities such as tachycardia or arrhythmias.

The successful virally mediated optogenetic inhibition of efferent motor pathways we describe here completes a quartet of proof-of-concept experiments for peripheral optogenetic neuromodulation – it is now possible to use translationally relevant approaches to optogenetically *excite* as well as *inhibit* both *afferent* sensory neurons and *efferent* motor neurons. Translation of this neuromodulation quartet into the clinic will require careful choice of initial disease targets, improvements to the viral vectors used, and greater understanding of the consequences of long-term opsin expression.

## Acknowledgements

The authors would like to thank A. Christensen, K.L. Montgomery, and members of the Delp and Deisseroth labs for insightful discussions and help with experiments.

## Supporting Information

**S1 Table. Selected virally mediated attempts.** Efficient transfection and function of an inhibitory opsin was difficult. Over 23 different AAV serotype, promoter, opsin, and fluorescent tag combinations were used before AAV6-hSyn-iC++-eYFP was chosen as the most effective for peripheral motor neuron inhibition. Adeno-associated virus (AAV), recombinant (r), self-complementary (sc).

**S1 Fig. Schematic of experimental force setup.** Motor axons of the sciatic nerve were stimulated electrically with a stimulation cuff placed around the proximal sciatic nerve of an anaesthetized mouse. The sciatic nerve was then illuminated with a blue laser light (473 nm) at the cuff. Fine-wire EMG electrodes recorded electrical activity of the medial gastrocnemius and a force transducer attached at the Achilles tendon recorded the contractile force of the lower limb muscles.

## References

1. Katz RT, Rymer WZ. Spastic hypertonia: mechanisms and measurement. Archives of physical medicine and rehabilitation. 1989;70(2):144–55. PubMed PMID: 2644919.

2. Sanger TD. Hypertonia in children: how and when to treat. Curr Treat Options Neurol. 2005;7(6):427–39. PubMed PMID: 16221366.

3. Gomez-Amaya SM, Barbe MF, de Groat WC, Brown JM, Tuite GF, Corcos J, et al. Neural reconstruction methods of restoring bladder function. Nat Rev Urol. 2015;12(2):100–18. doi:10.1038/nrurol.2015.4. PubMed PMID: 25666987; PubMed Central PMCID: PMC4324509.

4. Potter PJ. Disordered control of the urinary bladder after human spinal cord injury: what are the problems? Prog Brain Res. 2006;152:51–7. doi:10.1016/S0079-6123(05)52004-1. PubMed PMID: 16198693.

5. Wright EF. Medial pterygoid trismus (myospasm) following inferior alveolar nerve block: case report and literature review. General dentistry. 2011;59(1):64–7. PubMed PMID: 21613042.

6. Smyth MD, Peacock WJ. The surgical treatment of spasticity. Muscle & nerve. 2000;23(2):153–63. PubMed PMID: 10639605.

7. Stetkarova I, Yablon SA, Kofler M, Stokic DS. Procedure- and device-related complications of intrathecal baclofen administration for management of adult muscle hypertonia: a review. Neurorehabil Neural Repair. 2010;24(7):609–19. doi:10.1177/1545968310363585. PubMed PMID: 20233964.

8. Lowe PL, Lowe NJ. Botulinum toxin type B: pH change reduces injection pain, retains efficacy. Dermatol Surg. 2014;40(12):1328–33. doi:10.1097/DSS.0000000000000178. PubMed PMID: 25350125.

9. Hassel B. Tetanus: Pathophysiology, Treatment, and the Possibility of Using Botulinum Toxin against Tetanus-Induced Rigidity and Spasms. Toxins. 2013;5(1):73–83. doi:10.3390/toxins5010073. PubMed PMID: WOS:000315407200007.

10. Pollmann W, Feneberg W. Current management of pain associated with multiple sclerosis. CNS drugs. 2008;22(4):291–324. PubMed PMID: 18336059.

11. Deisseroth K. Optogenetics: 10 years of microbial opsins in neuroscience. Nat Neurosci. 2015;18(9):1213–25. doi:10.1038/nn.4091. PubMed PMID: 26308982; PubMed Central PMCID: PMC4790845.

12. Lin SC, Brown RE, Hussain Shuler MG, Petersen CC, Kepecs A. Optogenetic Dissection of the Basal Forebrain Neuromodulatory Control of Cortical Activation, Plasticity, and Cognition. The Journal of neuroscience: the official journal of the Society for Neuroscience. 2015;35(41):13896–903. doi:10.1523/JNEUROSCI.2590-15.2015. PubMed PMID: 26468190; PubMed Central PMCID: PMC4604228.

13. Robinson MJ, Warlow SM, Berridge KC. Optogenetic excitation of central amygdala amplifies and narrows incentive motivation to pursue one reward above another. The Journal of neuroscience: the official journal of the Society for Neuroscience. 2014;34(50):16567–80. doi:10.1523/JNEUROSCI.2013-14.2014. PubMed PMID: 25505310; PubMed Central PMCID: PMC4261087.

14. Zhang F, Prigge M, Beyriere F, Tsunoda SP, Mattis J, Yizhar O, et al. Red-shifted optogenetic excitation: a tool for fast neural control derived from Volvox carteri. Nature neuroscience. 2008;11(6):631–3. doi:10.1038/nn.2120. PubMed PMID: 18432196; PubMed Central PMCID: PMC2692303.

15. Bryson JB, Machado CB, Lieberam I, Greensmith L. Restoring motor function using optogenetics and neural engraftment. Current opinion in biotechnology. 2016;40:75–81. doi:10.1016/j.copbio.2016.02.016. PubMed PMID: 27016703.

16. Mahn M, Prigge M, Ron S, Levy R, Yizhar O. Biophysical constraints of optogenetic inhibition at presynaptic terminals. Nat Neurosci. 2016;19(4):554–6. doi:10.1038/nn.4266. PubMed PMID: 26950004.

17. Towne C, Montgomery KL, Iyer SM, Deisseroth K, Delp SL. Optogenetic control of targeted peripheral axons in freely moving animals. PloS One. 2013;8(8):e72691. doi:10.1371/journal.pone.0072691. PubMed PMID: 23991144; PubMed Central PMCID: PMC3749160.

18. Liske H, Towne C, Anikeeva P, Zhao S, Feng G, Deisseroth K, et al. Optical inhibition of motor nerve and muscle activity in vivo. Muscle & nerve. 2013;47(6):916–21. doi:10.1002/mus.23696. PubMed PMID: 23629741.

19. Bryson JB, Machado CB, Crossley M, Stevenson D, Bros-Facer V, Burrone J, et al. Optical control of muscle function by transplantation of stem cell-derived motor neurons in mice. Science. 2014;344(6179):94–7. doi:10.1126/science.1248523. PubMed PMID: 24700859.

20. Berndt A, Lee SY, Wietek J, Ramakrishnan C, Steinberg EE, Rashid AJ, et al. Structural foundations of optogenetics: Determinants of channelrhodopsin ion selectivity. Proceedings of the National Academy of Sciences of the United States of America. 2015. doi:10.1073/pnas.1523341113. PubMed PMID: 26699459.

21. Hentschke H, Stuttgen MC. Computation of measures of effect size for neuroscience data sets. The European journal of neuroscience. 2011;34(12):1887–94. doi:10.1111/j.1460-9568.2011.07902.x. PubMed PMID: 22082031.

22. Iyer SM, Montgomery KL, Towne C, Lee SY, Ramakrishnan C, Deisseroth K, et al. Virally mediated optogenetic excitation and inhibition of pain in freely moving nontransgenic mice. Nature biotechnology. 2014;32(3):274–8. doi:10.1038/nbt.2834. PubMed PMID: 24531797; PubMed Central PMCID: PMC3988230.

23. Ferreira V, Petry H, Salmon F. Immune Responses to AAV-Vectors, the Glybera Example from Bench to Bedside. Frontiers in immunology. 2014;5:82. doi:10.3389/fimmu.2014.00082. PubMed PMID: 24624131; PubMed Central PMCID: PMC3939780.

24. Arruda VR, Favaro P, Finn JD. Strategies to modulate immune responses: a new frontier for gene therapy. Molecular therapy: the journal of the American Society of Gene Therapy. 2009;17(9):1492–503. doi:10.1038/mt.2009.150. PubMed PMID: 19584819; PubMed Central PMCID: PMC2835266.

25. Govorunova EG, Sineshchekov OA, Janz R, Liu X, Spudich JL. NEUROSCIENCE. Natural light-gated anion channels: A family of microbial rhodopsins for advanced optogenetics. Science. 2015;349(6248):647–50. doi:10.1126/science.aaa7484. PubMed PMID: 26113638.

26. Montgomery KL, Yeh AJ, Ho JS, Tsao V, Mohan Iyer S, Grosenick L, et al. Wirelessly powered, fully internal optogenetics for brain, spinal and peripheral circuits in mice. Nature methods. 2015;12(10):969–74. doi:10.1038/nmeth.3536. PubMed PMID: 26280330.

27. Glickman S, Kamm MA. Bowel dysfunction in spinal-cord-injury patients. Lancet. 1996;347(9016):1651–3. PubMed PMID: 8642958.

28. Nakipoglu GF, Kaya AZ, Orhan G, Tezen O, Tunc H, Ozgirgin N, et al. Urinary dysfunction in multiple sclerosis. J Clin Neurosci. 2009;16(10):1321–4. doi:10.1016/j.jocn.2008.12.012. PubMed PMID: 19560927.

29. Stoffel JT. Detrusor sphincter dyssynergia: a review of physiology, diagnosis, and treatment strategies. Transl Androl Urol. 2016;5(1):127–35. doi:10.3978/j.issn.2223-4683.2016.01.08. PubMed PMID: 26904418; PubMed Central PMCID: PMC4739973.

30. Nussinovitch U, Gepstein L. Optogenetics for suppression of cardiac electrical activity in human and rat cardiomyocyte cultures. Neurophotonics. 2015;2(3):031204. doi:10.1117/1.NPh.2.3.031204. PubMed PMID: 26158013; PubMed Central PMCID: PMC4478752.

31. Nussinovitch U, Shinnawi R, Gepstein L. Modulation of cardiac tissue electrophysiological properties with light-sensitive proteins. Cardiovascular research. 2014;102(1):176–87. doi:10.1093/cvr/cvu037. PubMed PMID: 24518144.

